# Linking geography and mutation profiles across goat species

**DOI:** 10.64898/2026.07.09.737241

**Authors:** A. Bionda, P. Crepaldi, J. Prendergast, M. Naupane, M. Amills, B.J. Rosen, G. Tosser-Klopp, M. Milanesi, A. Talenti, The VarGoats Consortium

## Abstract

Recent studies have characterised the mutational profile across multiple mammalian species, highlighting substantial differences across lineages. However, none of these studies investigated whether mutation profiles and geography are significantly correlated. In this study, we present a multi-genome alignment spanning several *Capra* taxa, reconstruct the ancestral genome of *Capra hircus* and use it to characterize the mutational profiles across multiple *Capra* species by using the 1000 genomes VarGoats dataset. Results confirmed that the scale of differences among *Capra* species largely reflects their phylogenetic relationships, in particular with the Bezoar being genetically closer to domestic goats than to other wild species. Subsequently, we correlated the mutational profile and the geographical origin of the different individuals. In particular, ACG>ATG changes have the strongest correlation with longitude (r = –0.79, P-value = 3.02*10^-204^), while TCA>TGA are strongly correlated with latitude (r = −0.51, P-value = 4.30*10^-63^). We highlight how sequential dinucleotide mutations (SDMs) place cosmopolitan breeds closer to the sampling location, rather than the country of origin, showing how the recent relocation of cosmopolitan breeds to new continents is reshaping the genome of these animals. Finally, we used the mutational profile to predict the coordinate of origin of each animal in the dataset. In conclusion, we show the important role that geography had in shaping the genomes of domestic goats.

## Introduction

Mutation is a key process underlying the creation of new genetic variation and the emergence of novel species, populations and phenotypes (Acuna-Hidalgo et al. 2016), and is mainly caused by the combination of DNA replication errors and chemical and physical damage (Lindahl and Wood 1999; Chatterjee and Walker 2017). Mutations can also compromise cellular functions and reduce individual fitness, with particularly well-studied consequences in cancer and other diseases (Sharma et al. 2020). To limit the accumulation of such deleterious genomic alterations, mammalian species have evolved multiple families of genes dedicated to repairing DNA, which are essential for preserving genome integrity (L. Li et al. 2021). DNA repair mechanisms are complex, involving hundreds of genes acting coordinately across multiple pathways (L. Li et al. 2021).

Whilst DNA repair genes are fundamental to avoid the accumulation of mutations in genomes, they can be affected by mutations themselves, compromising their functional properties, and thus changing their ability to repair DNA damage. Most changes affecting DNA repair pathways have moderate or low impact on gene functionality (Zhou et al. 2013). However, even small variations in the efficiency of DNA-repair genes can cause changes to accumulate across genes and generations, potentially leading to significant differences in the pattern and frequencies of the different mutation types.

The patterns and frequencies of different mutation types across genomes of single individuals are globally referred to as the mutation spectrum. The somatic mutation spectrum has been characterized in several types of cancer, with databases such as COSMIC linking certain mutational profiles with specific DNA repair genes (Tate et al. 2019). Conversely, germline mutational spectra have been less studied. Evolutionary history, domestication events and selection pressures all contribute to the different mutational profiles of animal species, and complicate the analysis and investigation of the genes leading to imbalanced accumulation of mutations. Despite this, few studies have documented the mutation profiles across species. Harris and Pritchard characterized the spectrum of mutation across human populations, revealing that Europeans show an enrichment in TCC>TTC mutation (Harris and Pritchard 2017). The authors also highlighted how different primates have different mutational patterns (Harris and Pritchard 2017). More recent studies have shown how the germline mutation spectrum varies greatly across species (Beichman et al. 2023; Bergeron et al. 2023; Talenti et al. 2025). Bergeron et al. (2023) identified generation time and fertility among the traits linked to differences in the number of germline mutations per generation between vertebrate species. Beichman et al. (2023) characterized the mutation spectrum across mammalian species, with a particular focus on biased gene conversion (BGC), i.e., the process driving the increase in frequency of A/T to G/C, while decreasing G/C>A/T, likely be a major driver of the observed individual differences in mutation spectra.

Mutational profiles in livestock have been only recently characterized. Talenti et al. (2025) focused particularly on domesticated species, including cattle, pig and swamp buffalo. This work highlighted how mutational spectra across species partly reflects phylogeny, with closely related species sharing similar mutational profiles, but suggesting additional influences on mutational processes. It also identified mutational profiles specific for some species, such as the unusually low rates of C>T mutations in CpG contexts in pigs, and those shared among them, such as the TCC>TTC change, already identified by Beichman et al. (2023) as elevated not only in European humans but also in East Asian Indicine cattle. As part of their work, the authors released a Nextflow workflow called nSPECTRa that produces the mutational profiles for any dataset of any species for both individual single-nucleotide variants (SNV) and sequential dinucleotide mutations (SDM, i.e. when a second SNV change occurs directly after a first one) (Prendergast et al. 2019; Talenti et al. 2025).

Goats (*Capra hircus*) were among the first animal species to be domesticated, and subsequently spread throughout the world, and adapted to a variety of environments and management conditions (Zeder and Hesse 2000; J.A. Erven et al. 2025). Unlike other domesticated species, bezoar (*Capra aegagrus*), the ancestor of domestic goats, is still present in the Middle East, Caucasus and Anatolia. In addition to the bezoar, several other wild *Capra* species are still present in Europe, Africa and Asia, including the Markhor (*Capra falconeri*), Turs and multiple Ibex species. This makes goats an excellent example for studying within- and between-species divergence linked to drift, selection and mutation. To this purpose, the VarGoats consortium has generated a dataset comprising 1,372 genomes from various domestic breeds and their wild relatives, offering the scientific community a unique opportunity (Denoyelle et al. 2021; Lazzari et al. 2025).

In this study we use the dataset provided by the VarGoats consortium (https://www.goatgenome.org/vargoats.html) to explore the mutation spectra for the domestic goat (*Capra hircus*) and its closely related wild species (Lazzari et al. 2025). We focus particularly on investigating the link between geography and mutational signatures, showing the impact that breeding and adaptation has had on the genome of this important livestock species.

## Results and Discussion

In this work we characterized the mutational profiles of the *Capra* genus, with a particular focus on the link between individual changes and geography. The analysis of mutational profiles relies on the availability of accurate ancestral alleles for each variant so that the direction of base changes can be determined. We therefore generated a reference-free, multiple genome alignment of five domestic and one wild goat genome (*C. sibirica*; GCA_003182615.2), as well as 4 different outgroups (sheep, pig, human, and mouse genomes), using CACTUS (Armstrong et al. 2020). This multi-genome alignment could be of use for future studies involving any species of the *Capra* genus, making it a valuable resource for the scientific community. In addition to aligning genomes, CACTUS is capable of computing the ancestral genome at each inner node of the guide tree, albeit fragmented and not reflecting the chromosomal structure of the original genome. We therefore proceeded to reconstruct the ancestral state of the *Capra hircus* genome by using the alignment information and the ancestral genome at the appropriate inner node using nSPECTRa (Talenti 2025; Talenti et al. 2025).

This ancestral genome covers 89.8% of the total genome length (2,624,162,033 out of 2,922,813,246 bases), offering a good representation of the ancestral sequence of *Capra hircus*. While this manuscript focuses on the autosomes, the ancestral genome contains all the sequences from the original ARS1 reference genome, including the mitochondria (NC_005044.2), and the two (NW_017189516.1 and NW_017189517.1) and eleven (NW_017189563.1, NW_017189610.1, NW_017189618.1, NW_017189628.1, NW_017189685.1, NW_017189696.1, NW_017189885.1, NW_017189985.1, NW_017190040.1, NW_017190154.1, NW_017195709.1) scaffolds composing the X and Y chromosomes, respectively. This allowed us to accurately identify the derived allele for many variants in the VarGoats dataset (15,876,409/26,858,361; 59.11%), and therefore to offer a representation of the mutational profile of the species, both as SNV and SDM changes.

### The mutation profile of the *Capra* genus

We computed SNV and SDM mutation profiles using the nSPECTRa workflow. We initially compared the mutation profiles of all unrelated samples (UNREL; n. 1,087) by generating a principal component analysis (PCA) from the SNV and SDM changes (Figure 1). This analysis confirmed the marked genetic differences of wild species compared with domestic goats. The first two PCs of the SNV profile (Figure 1a) discriminated among species, with the bezoar clustering with domestic goats and markhors taking an intermediate position. Domestic goats form a tight cluster, separate from the other *Capra* species (Figure 1a); when considering non-exported goats only (i.e. animals sampled from the geographical origin of the breed) it is possible to see the differences between the three major geographic clusters (subpane in Figure 1a). On the other hand, the SDM-based PCA (Figure 1b) mirrored the population structure observed from genotype data directly (Figure 1c), primarily clustering individuals by continent of origin, with wild goats grouped within the Asian cluster. Notably, the third and fourth PCs of the SDM data provided better discrimination among *Capra* species (Supplementary Figure S1).

**Figure 1.**
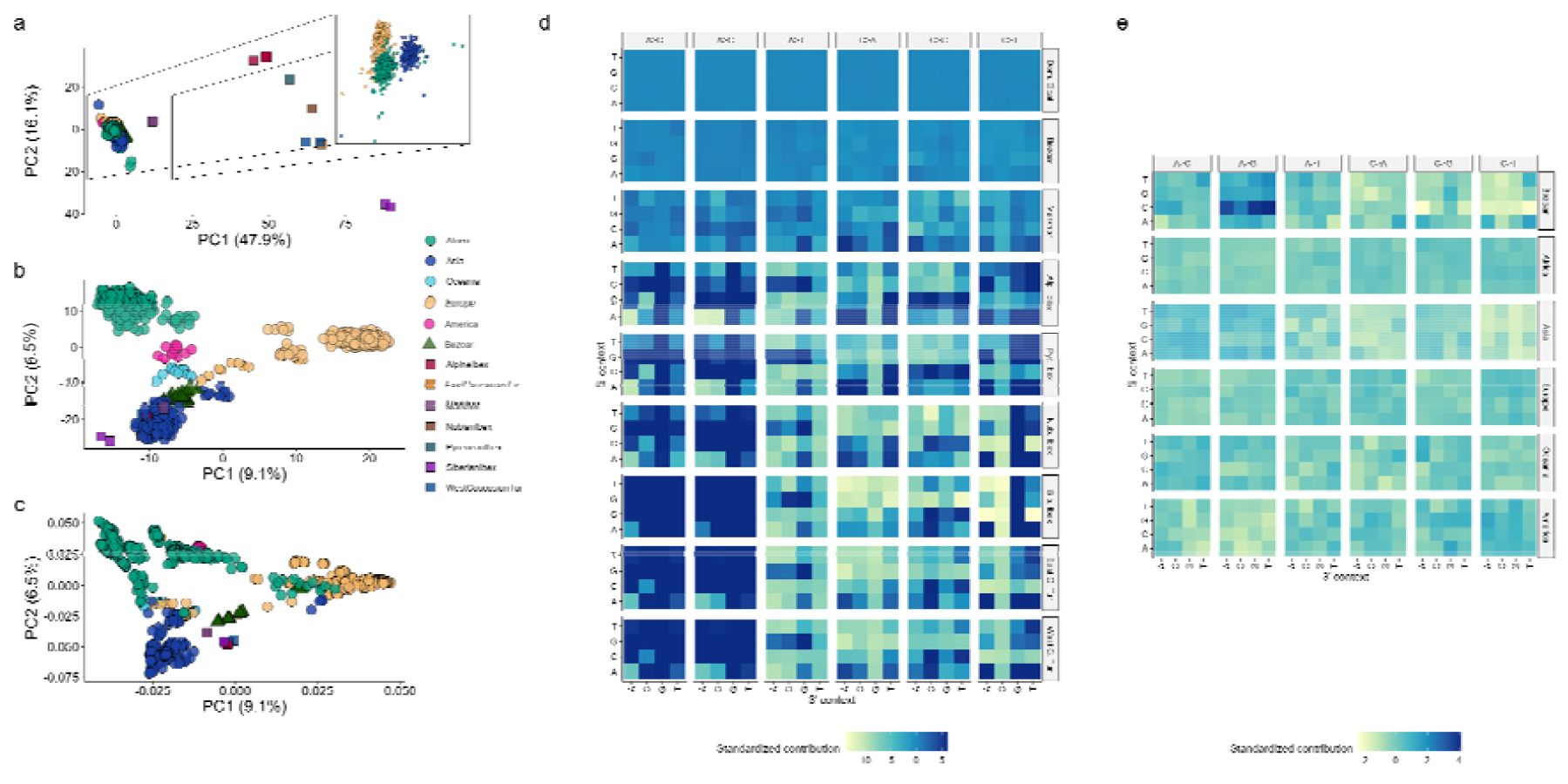
Cross-species comparison of the mutational profile. a) PCA computed on the SNV mutation profiles of the UNREL dataset, the zoomed section shows the SNV profile PCA on the non-exported domestic goats only (UNREL_NOEXPORTED); b) PCA computed from the SDM mutation profiles of the UNREL dataset; c) PCA computed from the genotypes of the UNREL dataset. d) Heatmaps of the SNV types of d) all the analysed Capra species (UNREL) and e) and domestic goats and bezoars only showing an increase in C[A>G]N changes (UNREL_BEZ).

The heatmap of the SNVs across species (Figure 1d) supported the PCA findings, confirming that the differences between domestic goats and bezoars were minimal when compared to the other wild species. Similarly, markhors exhibited intermediate mutation patterns between domestic goats and other wild species. Additionally, differences were observed among wild species: for example, both East and West Caucasian turs, along with the Siberian ibex, displayed similarly elevated frequencies of A>G transitions compared to the Alpine, Pyrenean, and Nubian ibexes.

When focusing specifically on domestic goats and bezoars (UNREL_BEZ), a clear separation emerged in both PCA (Supplementary Figure S2) and heatmap analyses, with evidence that the mutation profile of bezoars more closely resembled that of domestic goats from Asia. Specifically, the heatmap revealed that bezoars had a higher frequency of C[A>G]N changes and lower frequencies of C[C>T]N and C[C>G]A changes compared to domestic goats (Figure 1e).

We then identified which changes better discriminated the different species by training and applying an eXtreme Gradient Boosting (XGBoost) model to our SNV mutation profile data (Chen and Guestrin 2016). This method has previously been described as an effective approach to both classify individuals according to their mutational profile, and to select which features are better at discriminating each class (Talenti et al. 2025). The model trained on UNREL successfully classified individuals as domestic goats or bezoars, but fails to discriminate between the different wild goat species due to the reduced sample sizes (Supplementary Figure S3). Similarly, it fails to identify changes able to discriminate between different species, with the exception of the bezoar, for which the C[A>G]G was identified as the most differentiating change (Figure 2a and 2b), consistent with the results in Figure 1e.

**Figure 2.**
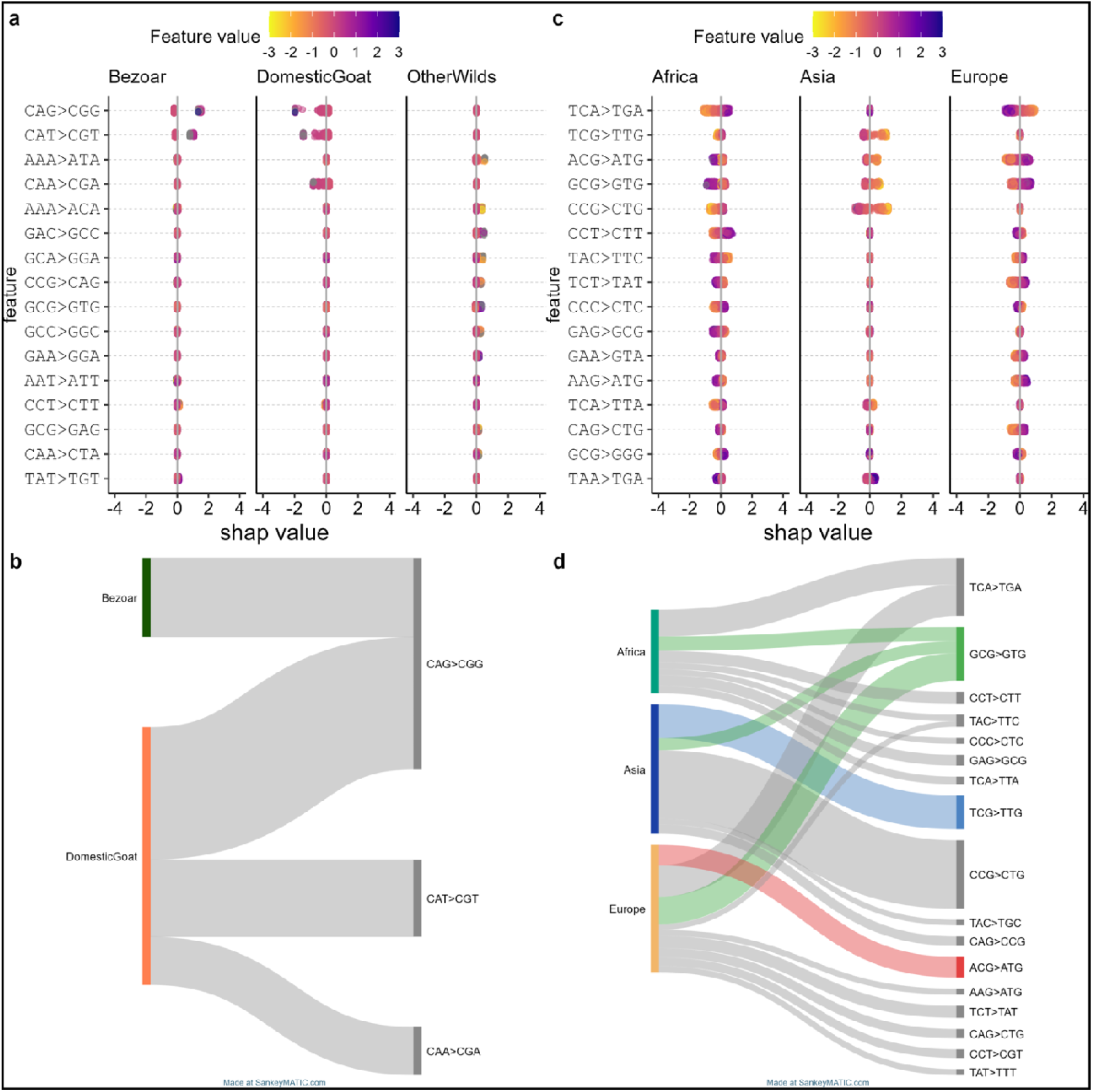
SHAP values (a) and Sankey diagram (b) of the SNV that best discriminate Capra hircus, Capra aegagrus (i.e. Domestic goat and bezoar), and all the other available Capra species grouped together (UNREL dataset); and SHAP values (c) and Sankey diagram (d) of the SNV that best discriminate domestic goats from different continents (UNREL_NOEXPORTED dataset). In a and c, the x axis indicates the impact of the change reported on the y-axis, with positive values associated with enrichment and negative with depletion of the related mutation. Specifically, more spread SHAP values for a species reflect a stronger ability of the change to discriminate it from the others. Points are coloured according to the mutation value. In b and d, the SNV values with a variance in the SHAP values for the corresponding mutation > 0.05 for any species are reported. A higher variance reflects a stronger association between a change and a species.

### The mutation profile of the *Capra hircus*

The second part of this study focused exclusively on the mutation profile of domestic goats. Specifically, we initially analysed the UNREL_NOEXPORTED dataset, excluding animals sampled on continents different from where the breed was derived, to better explore geographic patterns shaped by long-term selection. The SNV-based PCA revealed a clear separation of individuals by continent of origin: Asian goats were isolated along PC1, which accounted for 24.5% of the variance, while African and European goats were separated along PC2, accounting for 6.1% of the variance (Figure 1d). Minimal overlaps were observed, which can be explained by breed history or current geographic locations. For example, the Norwegian breed reared in Tanzania was originally imported from Northern Europe (Msalya et al. 2017), or the Palmera goats are formally Spanish, but arrived in the Canary islands prior to the Spanish colonization and have African ancestry (J.A.M. Erven et al. 2025).

When analysing the SDM-based PCA, the continental separation along the first two PC was even more distinct than that obtained using SNV or genotype-based PCA, with PC1 (11.5% of variance) separating European goats and PC2 (7.9% of variance) discriminating between Africa and Asia (Supplementary Figure S4). Similarly to the SNV results, only a few exceptions were identified, such as the Russian samples intermediate between Asian and European clusters; the African Burundi breed reared in Switzerland; and the Palmera goats, clustered with European breeds but shifted toward the African group. Conversely, the Norwegian breed from Tanzania grouped with African goats, though with a shift toward Europe.

An XGBoost model was used to predict the continent of origin of non-exported domestic goats based on both SNV and SDM profiles, achieving accuracies of 96% and 100%, respectively. SHAP values were computed for both models to identify the most informative features. These analyses revealed that, whilst most SNV variants distinguishing goats from different continents were largely unique to each class (Figure 2c and 2d), few were able to discriminate between multiple continents, with GCG>GTC substitutions being able to distinguish individuals from all three examined continents. A notable finding is the TCG>TTG change, which is able to exclusively identify Asian goats. Noticeably, this group showed an overall significant depletion in C>T substitutions in CpG contexts (Kruskal-Wallis adjusted P-value = 3.5 * 10^-49^ and 7.9 * 10^-133^ for Africa-Asia and Europe-Asia comparison; Supplementary Figure S5); therefore with TCG>TTG in particular emerging as the most discriminating change. A reduction in CpG-context deamination has already been observed in Asian pigs, where an even stronger decrease in the same class of changes was reported (Talenti et al. 2025), suggesting an intriguing geographical parallel with goats. Further analyses will focus on performing a cross-species evaluation of the variants impacting the homologous repair machinery and the mismatch repair mechanisms. Additionally, Asian goats displayed a marked enrichment in A>C and A>G substitutions, particularly TAC>TGC and CAG>CCG.

### Sequential dinucleotide mutations correlate to geographical patterns in *Capra hircus*

A more in-depth analysis of the SDM-based PCA revealed that components greater than PC2 were able to differentiate goat populations at the country level within each continent. For example, PC3 clustered goats along the North-South African axis, with breeds from Northern Africa (Tunisia, Morocco, Mali), Equatorial Africa (Uganda, Tanzania, Kenya), and Southeastern Africa (Zimbabwe, Mozambique, Malawi and Madagascar) clearly separated in three groups (Figure 3a). An interesting observation is that PC4 distinctly separated goats from Madagascar, regardless of breed and even when those breeds were also present in other African countries (Figure 3a). A similar pattern was observed in the genotype-based PCA (Supplementary Figure S6a), and this is probably the result of extensive founder effects experienced by these populations led by the insularity (Cardoso et al. 2018). What is interesting is that this genetic uniqueness can be seen also at the mutational profile level, with a shift in the mutational spectrum that differs from that observed in the African continent. PC6 separated goats from different Asian countries along an East-West axis, with Korean and Chinese goats on one end of the axis, and Pakistani and Iranian goats on the other (Figure 3b).

**Figure 3.**
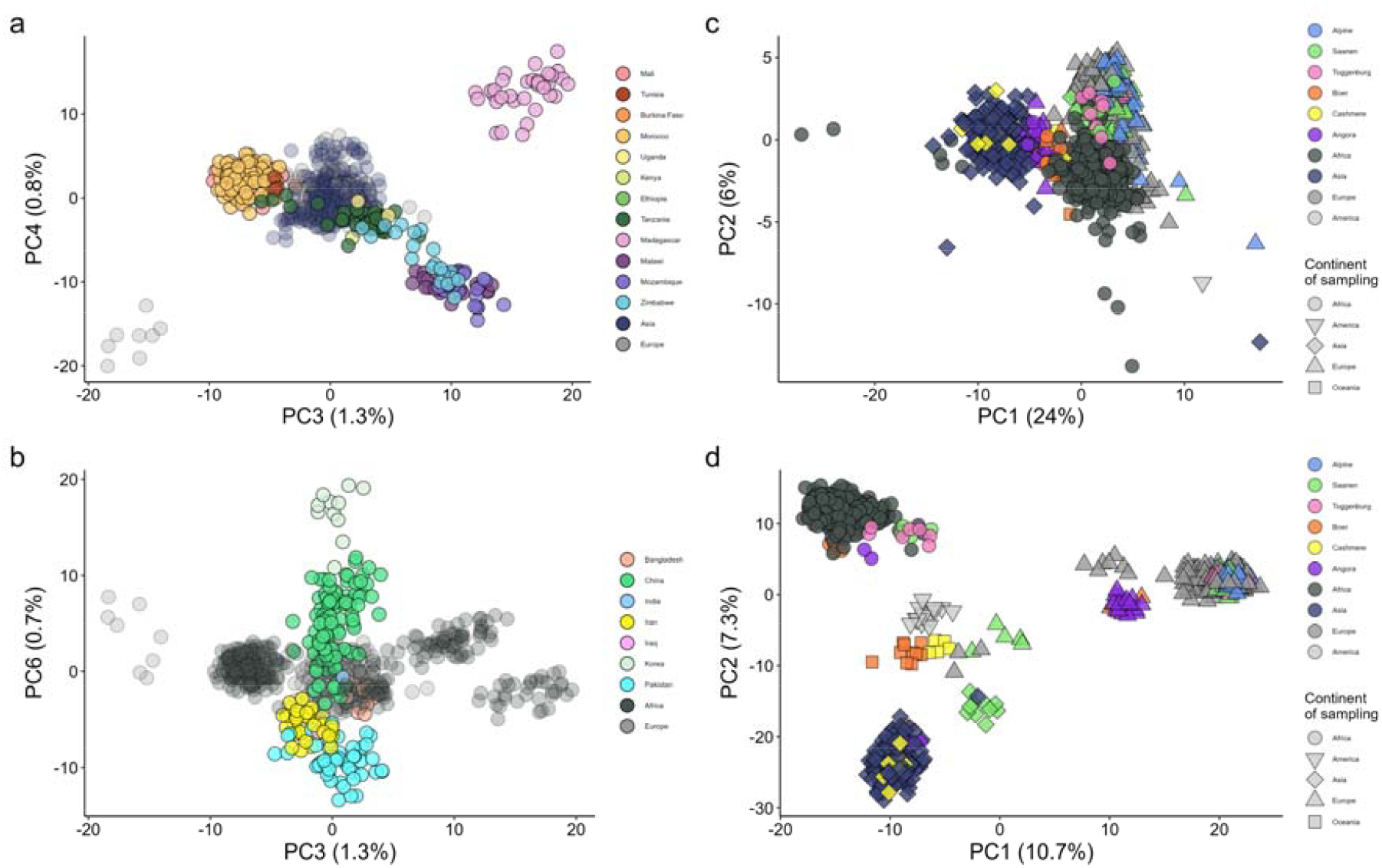
Domestic goat mutational profile, computed on the SDM profile of the UNREL_NOEXPORTED dataset differentiating African goats along PC3 and 4 (a) and Asian goats along PC3 and 6 (b). PCA of UNREL_DOMESTIC dataset, with a focus on transboundary breeds, showing that they cluster within their continent of origin (Europe for Alpine, Saanen, and Toggenburg; Africa for Boer; Asia for Angora and Cashmere) according to SNV profile (c), and within the continent of sampling according to SDM (d).

Further interesting considerations come from the analysis of transboundary breeds. By analysing the SNV profile of the UNREL_DOMESTIC dataset (Figure 3c), we can see that these goats clustered with their continent of origin, similarly to what is observed with the genotyping data (Supplementary Figure S6b). For instance, Cashmere and Angora cluster with the remaining Asian goats, Boers are close to the African goats, and Alpine, Saanen, and Toggenburg cluster near their European counterparts. However, the pattern changes when SDM data are used, as the majority of goats from transboundary breeds are placed within, or clearly shifted towards, the continent where they are currently raised (Figure 3d). This specific pattern is possibly due to two distinct features of the SDMs: 1) their rarity when compared with SNV, and 2) the second change is usually a more recent change of an older haplotype (Prendergast et al. 2019). To assess whether this applies to our data as well, we extracted the first and second changes composing each SDM, and computed the mutational profiles on them separately. In fact, neither the mutational profiles of the first or second sites composing the SDMs identify the geographical effect on the exported goats (Supplementary Figure S7). Moreover, the first change highlights a population structure very similar to the SNV dataset (Supplementary Figure S7a), whereas the second change displays a different structure (Supplementary Figure S7b), further supporting the theory that the second change is more recent and happened on top of a previous one.

### Predicting the geographic origin of goats from the mutational profile

After detecting a relationship between continent of origin and the mutational profile of the domestic goat, we further investigated this relationship by calculating the pairwise correlation between geographic coordinates and the frequencies of all possible SNV changes. Then, we modeled the frequency of each substitution type as a function of longitude, latitude, and their interaction using a generalized linear regression model. These analyses were limited to a total of 943 non-exported individuals for which coordinates were available (Supplementary Table S1). The regression results showed R^2^ values ranging from 0.006 to 0.65. It is interesting how five out of the ten most correlated changes involve C>T substitutions in a CpG context (Supplementary Table S2). The suggestive link between deamination in CpG context and geography is further strengthened by the ACG>ATG change showing the strongest correlation with longitude (r = –0.79, FDR-corrected P-value = 3.02*10^−204^ ; Figure 4a). The strongest link with latitude was observed with the TCA>TGA change (r = –0.51, FDR-corrected P-value = 4.30*10^−63^ ; Figure 4b), further hinting at a specific link between deamination in CpG sites and the East-West gradient in the Eurasian continent (Figure 4c).

**Figure 4.**
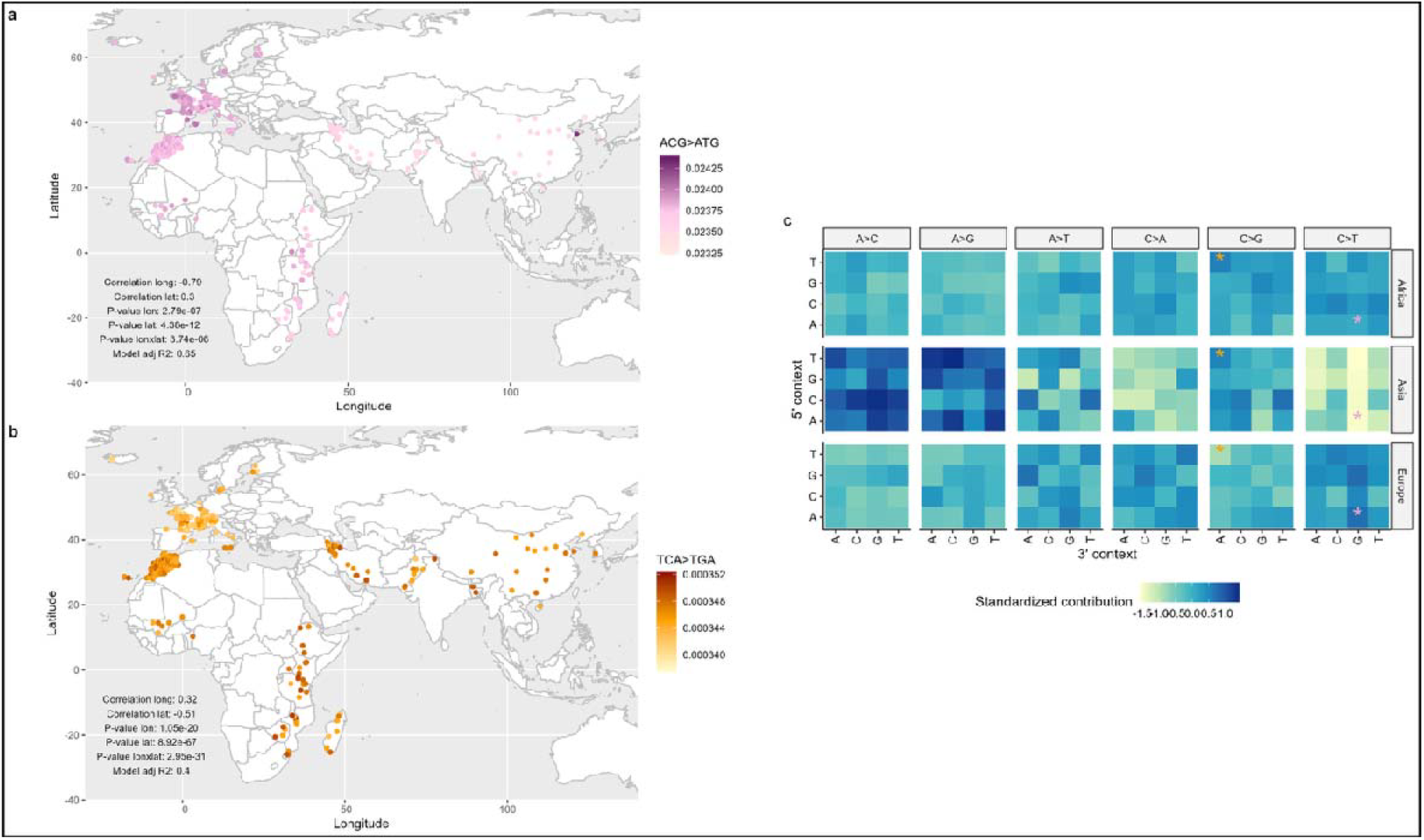
Correlations between geography and mutational changes in the domestic goat. Geographical distributions of normalized frequencies of the ACG>ATG (a) and TCA>TGA (b) changes, showing a significant correlation with longitude and latitude, respectively; the P-values for longitude, latitude, and their interaction, as well as the whole model R^2^, are also indicated. (c) Heatmap of the different normalized frequencies of SNV mutations in the different geographical groups, with the ACG>ATG change marked in pink and TCA>TGA marked in orange, confirming the East-to-West and North-to-South correlations for the two changes, respectively. Analyses were performed on the UNREL_NOEXPORTED dataset.

To identify the best predictor of the geographic origin of the investigated domestic goats, we implemented a Gradient Boosting Machine (GBM) model to infer sample longitude and latitude using SNV and SDM mutation profiles, their respective PCs, or the genotype-based PCs. Longitude was consistently better predicted than latitude across all models (Table 1). The best performance was achieved using the genotype-based PCs, closely followed by the SDM-based PCA, with very similar statistics. The mean Haversine distance between the observed and predicted geographic locations was 301 km for the model trained on genotype PCs and 461 km when trained on the SDM PCs, while the latter outperforms the former in predicting the latitude of origin. Importantly, although the genotype-based model is more precise in predicting the coordinates of origin, it is uninformative of the underlying biology. On the other hand, the SDM and SNV prediction models are linked to differences in the DNA mutation/repair mechanisms within the goat species. This offers an important insight into how population divergence over a short period of time led to differences in their response to DNA damage or mutational pressure, and therefore to preferential accumulation of specific changes.

**Table 1.** Statistics on the GMB models for predicting latitude and longitude of the test dataset composed of non-exported domestic goats (UNREL_NOEXPORTED). All values are reported as mean ± standard deviation across the five runs of each model.

| Model | N. levels | Longitude |  |  | Latitude |  |  | Haversine distance |  |
| --- | --- | --- | --- | --- | --- | --- | --- | --- | --- |
|  |  | RMSE | R<sup>2</sup> | MAE | RMSE | R<sup>2</sup> | MAE | Median (km) | Mean (km) |
| SNV | 96 | 12.67 ± 0.48 | 0.89 ± 0.01 | 8.56 ± 0.38 | 10.39 ± 0.31 | 0.76 ± 0.01 | 6.92 ± 0.2 | 883.42 ± 51.03 | 1274.77 ± 28.2 |
| SNV PCs | 96 | 12.29 ± 0.45 | 0.90 ± 0.01 | 8.25 ± 0.33 | 8.91 ± 0.19 | 0.83 ± 0.01 | 5.86 ± 0.1 | 794.22 ± 27.57 | 1180.9 ± 32.32 |
| SDM | 2288 | 8.68 ± 0.23 | 0.95 ± 0 | 6.19 ± 0.23 | 6.97 ± 0.09 | 0.90 ± 0 | 4.6 ± 0.14 | 668.75 ± 44.22 | 898.61 ± 20.15 |
| SDM PCs | 96 | 4.70 ± 0.17 | 0.99 ± 0 | 3.24 ± 0.08 | 3.35 ± 0.06 | 0.98 ± 0 | 2.31 ± 0.07 | 338.50 ± 13.12 | 461.02 ± 4.65 |
| Genotype PCs | 96 | 3.54 ± 0.27 | 0.99 ± 0 | 1.89 ± 0.15 | 4.01 ± 0.26 | 0.96 ± 0 | 1.84 ± 0.18 | 175.53 ± 14.05 | 300.68 ± 19.92 |

When the same models were used to predict the geographic locations of all domestic goats with available coordinates, including exported individuals (UNREL_DOMESTIC), the results were consistent with known breed origins. The genotype PCs model yielded the most accurate predictions, followed by the SDM PCs model. For example, all Boer and Creole goats were predicted to have originated from Africa, regardless of their sampling location. The Boer is one of the main and most spread meat goat breeds and was developed by Dutch breeders in South Africa, starting from local stocks likely crossed with animals of European and/or Indian origin (Ramsay et al. 2000). The Creole goat, instead, is raised in the Caribbean area, including Guadalupe Island, where sampling was carried out, but represent a mixed population with European (in particular from the Iberian region) and, to a greater extent, African origin (Naves et al. 2001; Sevane et al. 2018). European dairy breeds such as Saanen and Toggenburg were predicted along a trajectory pointing toward Western Europe, whereas Cashmere goats from Oceania were predicted to originate in Northern (genotype PCs) or North-Eastern (SDM PCs) Africa. Angora goats sampled in Europe or Africa were placed in northern Africa using the SDM PCs, or more accurately in the Middle East with the genotype PCs (Figures 5a and 5b). Notably, Angora goats sampled in Madagascar were predicted to be located on the same island, in agreement with other analyses. Models based on other predictors showed more dispersed predictions for goats of the same breed but generally produced similar overall patterns. The availability of geographical coordinates provided by the VarGoats project allows us to identify geographical patterns looking at each individual mutational profile, overcoming the limitation of summarising the profile across populations or continents (Figure 4c). However, it is to be noted that accurate coordinates were not available for all the samples and/or geographic regions; therefore, only 1,007 animals with individual- or district-level coordinates were considered. This particularly affected East Asian animals, where the country of origin was often the only spatial piece of information available, limiting the ability to predict the exact location of the several goats that originated there, including cosmopolitan breeds such as Cashmere and Angora.

**Figure 5.**
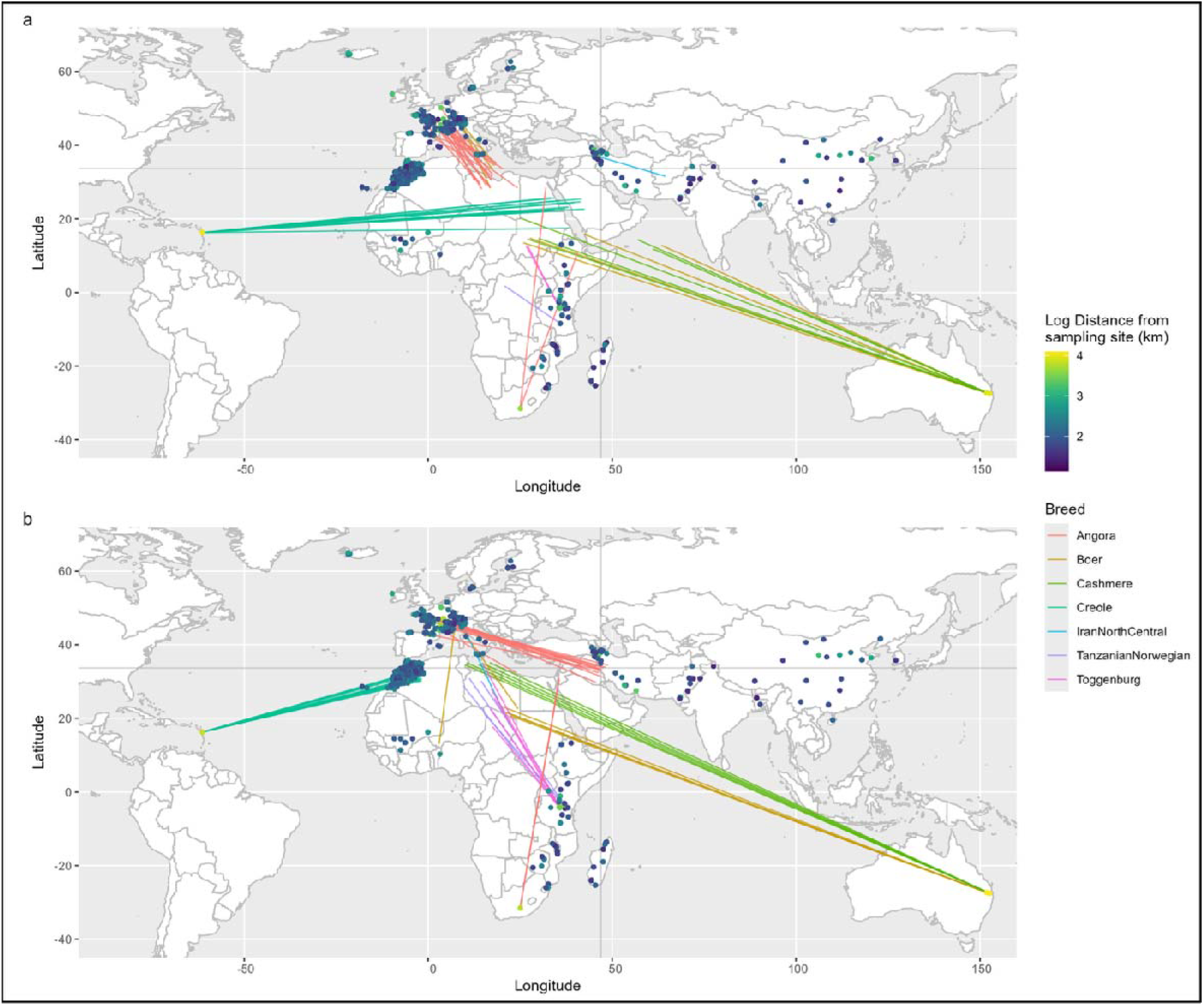
Predicted geographic locations of exported goats using the GBM models trained with SDM PCs (a) or genotype PCs (b) as predictors. All domestic goats with available geographic coordinates are shown at their sampling locations, represented as points coloured according to the Haversine distance between the observed and predicted locations. For goats with the top 5% highest distances, a line segment coloured by breed connects the observed and predicted coordinates (the latter calculated as the mean predicted location across all model runs).

While our work uncovered interesting patterns in the mutational profile of the *Capra* lineage, it still presents some limitations. As previously mentioned, the limited numbers of animals belonging to the different non-Bezoar wild *Capra* species is sufficient to highlight the relationships across them, but does not allow us to detect the more nuanced differences between them. Moreover, to achieve a fully unbiased representation of the mutational profile, individuals of each species should be aligned and variants called against their own species-specific reference genome, and all the individual references would be used to define a more comprehensive ancestral state of the *Capra* genus. However, at the time of the graph generation, this was not viable due to the limited number of high quality reference genomes for most of the species considered. Another previously noted limitation concerned the lack of high-resolution geolocalisations for some of the samples in the dataset. The VarGoats consortium variant database is a remarkable, albeit imperfect, resource with comprehensive metadata available for most of its animals, though not for all of them. As a consequence, some parts of the world could not be included in the geographical predictions. Finally, this study focuses exclusively on the domestic and wild goats, without placing them in the context of more distant species. Future studies will address this limitation by analysing these individuals in the context with other species, including other Bovidae and domesticated animals, allowing us to further understand the diversity and uniqueness of goat species.

## Conclusions

In this study we generated a multiple genome alignment involving multiple *Capra* species, a high-quality *Capra hircus* ancestral genome, and the characterization of the mutational profile in multiple *Capra* species. We found a clear link between geography and the *Capra hircus* mutational profile, and in particular how SDMs cluster transboundary animals in their sampling location, unlike the SNV changes that place them in their country or continent of origin. The link between the mutational profile and geography is strong enough to allow us to predict the coordinates of origin of the animals analysed directly from the spectrum with good accuracy. In conclusion, we demonstrate how short-term geographical adaptation can lead to measurable changes in the rate of mutation of the genome of domestic goats.

## Materials and methods

### Defining the ancestral genome

Germline variants called from whole genome resequencing are representative of the current genomic landscape of a given species, being the product of evolutionary forces such as mutation, selection and drift. In order to compare the mutational profile across any type of groups of individuals (e.g. by continent), it is critical to establish the allele at each position in the genome prior to the lineages that are studied having split. To achieve this, we generated a multiple genome alignment database composed of 10 genome assemblies (Table 2) using CACTUS (v2.0.4) (Armstrong et al. 2020).

**Table 2.** Genomes included in the CACTUS multiple genome alignment.

| Accession | Tree ID | Species | Breed | Length | Number of Sequences | Reference |
| --- | --- | --- | --- | --- | --- | --- |
| GCA_004361675.1 | BengalBlackGoat | Capra hircus | Bengal Black goat | 3041623660 | 3972 | (Siddiki et al. 2019) |
| GCA_009823495.1 | PashminaCashmereGoat | Capra hircus | Cashmere goat | 2941702774 | 48370 |  |
| GCA_015443085.1 | SaanenGoat | Capra hircus | Saanen goat | 2696218280 | 1362 | (R. Li et al. 2021) |
| GCF_001704415.1 | SanClementeGoat | Capra hircus | San Clemente goat | 2922813246 | 29907 | (Bickhart et al. 2017) |
| GCA_000317765.2 | YunnanBlack | Capra hircus | Yunnan Black goat | 2809418462 | 102895 | (Dong et al. 2013) |
| GCA_003182615.2 | CapraSibirica | Capra sibirica |  | 2733031394 | 85609 |  |
| GCF_016772045.1 | OvisAries | Ovis aries |  | 2628146905 | 142 | (Davenport et al. 2022) |
| GCF_000003025.6 | SusScrofa | Sus scrofa |  | 2501912388 | 613 | (Warr et al. 2020) |
| GRCh38 | HomoSapiens | Homo sapiens |  | 3099750718 | 194 |  |
| GRCm39 | MusMusculus | Mus |  | 2728222451 | 61 |  |

In order to run CACTUS, we first generated a phylogenetic guide tree detailing the relationships between the genomes present in the graph. To do so, we calculated pairwise distances using MASH (v2.2) (Ondov et al. 2016) with options `--k 32 --sketch_size 10000`, and then drew the tree using the UPGMA implementation in Biopython (Cock et al. 2009). The resulting phylogenetic tree was visualized using iTOL (v7.2, Figure 6; tree available on zenodo) (Letunic and Bork 2024). While constructing the alignments, CACTUS infers the ancestral state of each base in the genome at each split of the guide tree. These ancestral states are not reflective of the chromosomal structure of the genomes it refers to; instead, they are scattered across a set of individual contigs, aligned to each of the genome represented in the alignments.

**Figure 6.**
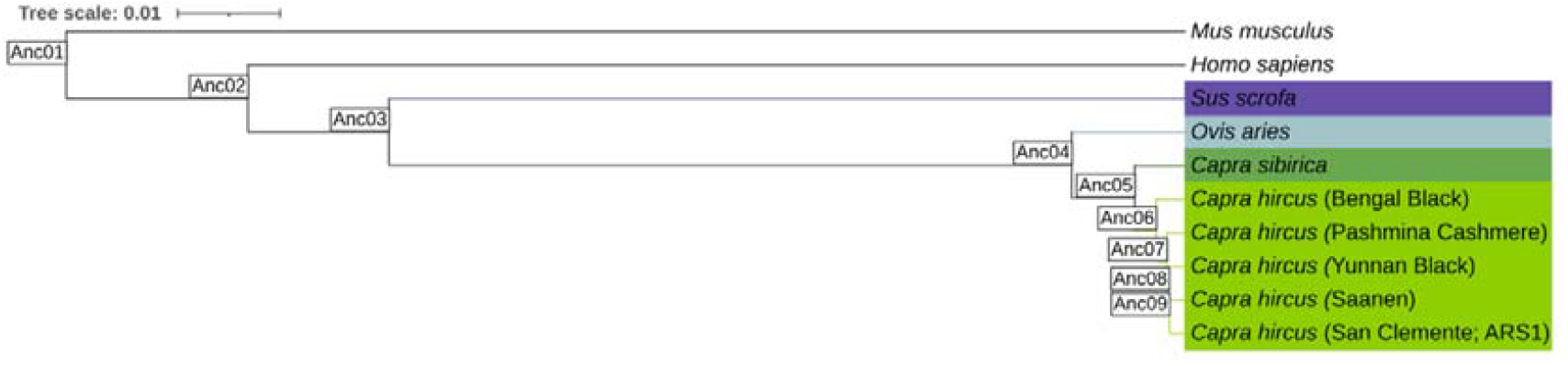
Phylogenetic tree derived from the mash distances between the genome assemblies used to generate the multiple genome alignment database of the capra species.

The nSPECTRa workflow (Talenti 2025; Talenti et al. 2025) takes the HAL alignments and anchors the ancestral state at each position in the desired reference genome, allowing for downstream analysis of the mutation profile. In this study we used Anc05 as the ancestral genome representative of the *Capra* genus.

### Dataset preparation

This study relies on the whole genome sequencing variant calls from the VarGoats project (Denoyelle et al. 2021), and consisted of 1,372 individuals, of which 1,327 were *Capra hircus* individuals and 45 were individuals of wild goat species (Supplementary Table S1). Genotypes have been filtered and imputed (Lazzari et al. 2025). Due to the mutational profile being representative of the broad differences between populations, we grouped individuals based on their continent of origin, as detailed.

The input dataset, after the exclusion of related individuals, comprised 1087 goats of 65 populations, sampled in Europe (N = 405), Africa (N = 398), Asia (N = 215), Oceania (N = 15), and America (N = 15), as well as 39 wild goats (N = 5 Alpine Ibex, N = 25 Bezoar, N = 1 East Caucasian Tur, N = 2 Markhor, N = 1 Nubian Ibex, N = 2 Siberian Ibex, N = 2 West Caucasian Tur). Related animals have been removed using KING (v2.2.7) (Manichaikul et al. 2010) and parameters `--unrelated --degree 2 -- sexchr 30`, removing 200 related individuals. Genotype-based PCA has been computed using plink (v1.9.0-b.8) (Chang et al. 2015) with options `--double-id --pca 96 --cow --maf 0.05 --geno 0.05` on a total of 8,756,382 variants and 1,087 individuals.

We then proceeded with the generation of three different datasets for the downstream analyses. The first dataset includes unrelated individuals for all *Capra* species (UNREL). The second includes unrelated individuals belonging to the domestic goats (*Capra hircus*) and bezoars (*Capra aegagrus*; UNREL_BEZ) species. The third dataset includes only the domestic goats (UNREL_DOMESTIC). The UNREL_DOMESTIC dataset was then filtered further to selectively retain goats sampled within the continent of origin: 1) Saanen, Toggenburg, and Alpine were excluded if not sampled in Western Europe, 2) Angora and Cashmere if not sampled in Asia, 3) Boers if not sampled in Africa, and 4) Creole goats from Guadeloupe were excluded because they were imported in recent times from Europe (UNREL_NOEXPORTED). Information on the composition of the different datasets can be found in Supplementary Table S1.

### Goat mutation profile

We then proceeded with the downstream analyses using nSPECTRa v1.1.0 for the ancestral genome generation, and v1.1.3 for the computation of the mutational profiles for both individual changes (SNV) and sequential dinucleotide mutations (SDM).

We have carried out the analyses on two distinct subsets: a dataset with all unrelated individuals, including animals of different species; and a dataset with only unrelated *Capra hircus* individuals (CHI, n. 1,048). Each dataset is processed in nSPECTRa independently.

We ran the workflow with the following options for UNREL: --reference SanClementeGoat --target Anc05 --mutyper --sdm --k 3,5,7 --chunk_size 1000000 -- max_derivate_allele_freq false --imputed --vep_container ‘ensemblorg/ensembl-vep:release_107.0’ --species capra_hircus. We ran the workflow for the CHI dataset using the same options, with the exception of setting `-- max_derivate_allele_freq 0.98`.

Raw counts have been normalized in two steps, as detailed in Talenti et al. (2025): 1) by the ancestral genome K-mer counts for the initial state; and 2) with the median of ratios. Briefly, we computed the geometric mean of each change type across all samples, then computed the ratio of each sample’s change counts to this reference. Finally, we normalized each sample with the median of these ratios for that individual. This method allows for a direct comparison of the mutation profiles for different individuals by effectively placing them on the same scale (Talenti et al. 2025). Individual changes composing the SDM were extracted using the bcftools view command (Danecek et al. 2021), and then annotated using mutyper (DeWitt et al. 2023).

We then visualized the normalized counts using the principal component analysis (PCA) function implemented in prcomp (stats R package). We generated heatmaps of the changes for each population using a modified version of the plot_standardized_profile_heatmap function from the MutationalPatterns R package (Manders et al. 2022).

To identify which mutation types best differentiate populations and species we fitted the normalised mutation spectra as features in multi-class extreme gradient boosting models using the xgbtree method in the caret R package with five-fold cross-validation (Kuhn 2008; Manders et al. 2022). To assess model performance 20% of the individuals were left out of the data prior to model training to act as an independent test set from which model metrics such as accuracy could be derived. To identify the sets of genetic changes that best differentiate populations we calculated SHAP values from these models using the xgboost R package (Lundberg and Lee 2017).

A gradient boosting machine (GBM) model was trained to classify samples from UNREL and UNREL_NOEXPORTED datasets into groups (e.g., species or continent) using the caret R package (Kuhn 2008) and xgboost model (Chen and Guestrin 2016). Specifically, the dataset was split into 80% training data and 20% test data, stratified according to the categories to be analysed. The xgbtree method was applied with five-fold cross-validation and the model’s performance was evaluated on test data by estimating accuracy, precision, recall, and F1-score, as well as by producing a confusion matrix. SHAP values were computed using the xgboost R package to interpret each change contribution to the groups’ differentiation.

### Geographical link analysis

Pairwise correlations of longitude and latitude with each possible SNV change were calculated using the cor.test function from the stats R package. P-values were adjusted for multiple testing via the Benjamini-Hochberg procedure, and significance was set at FDR-corrected P-value = 0.05.

Moreover, a generalized linear regression model was fitted with the glm function of the stats R package using each possible SNV as a response and latitude, longitude, and their interaction as fixed factors.

Lastly, the same GBM model presented in the previous section was applied to model longitude and latitude of each sample from: 1) SNV normalized mutation profiles; 2) eigenvector values obtained from the PCA performed on these SNV mutation profiles; 3) SDM normalized mutation profiles; 3) eigenvector values obtained from the PCA performed on the SDM profiles; 4) eigenvector values obtained from the PCA performed on genotyping SNP data. For SDM and genotyping SNP data, only the first 96 principal components were used, in order to make the analysis more comparable to the ones performed on SNVs.

The model was trained and tested on the UNREL_NONEXPORTED dataset individuals with available coordinates, splitting them using an 80/20 ratio. Each model was performed five times using different training/test data. The performance of the models was evaluated with the root-mean-square error (RMSE), the mean absolute error (MAE), and the R^2^ score. Each model was then used to predict the location of all the domestic goat samples, including the exported ones. The difference between the observed and the predicted location of the samples was estimated using Haversine distance (i.e., the shortest distance over Earth’s surface) by using the pracma R package’s haversine function (Borchers 2023).

## Data availability

The data underlying this article, including the multiple genome alignment file in HAL format, the ancestral genome, the mutation profiles and the K-mer counts are available in Zenodo (DOI: 10.5281/zenodo.20054870).

The VarGoats dataset is available on ENA under the umbrella project PRJEB37507, which includes the sequences generated for the project. Additional sequences were retrieved from public data. The 1,372 animals, including the publicly available ones, are listed on the VarGoats website (https://www.goatgenome.org/vargoats_data_access.html).

## Acknowledgements

AT is funded by the Jamieson Bequest fund at the School of Biodiversity, One-Health and Veterinary Medicine of the University of Glasgow. The list of members of the VarGoats consortium is available in the Appendix

## Notes

https://dx.doi.org/10.5281/zenodo.20054870

### Competing Interest Statement

The authors have declared no competing interest.

