## Supplementary figures and images for "Linking geography and mutation profiles across goat species"

### Supplementary Figure S1

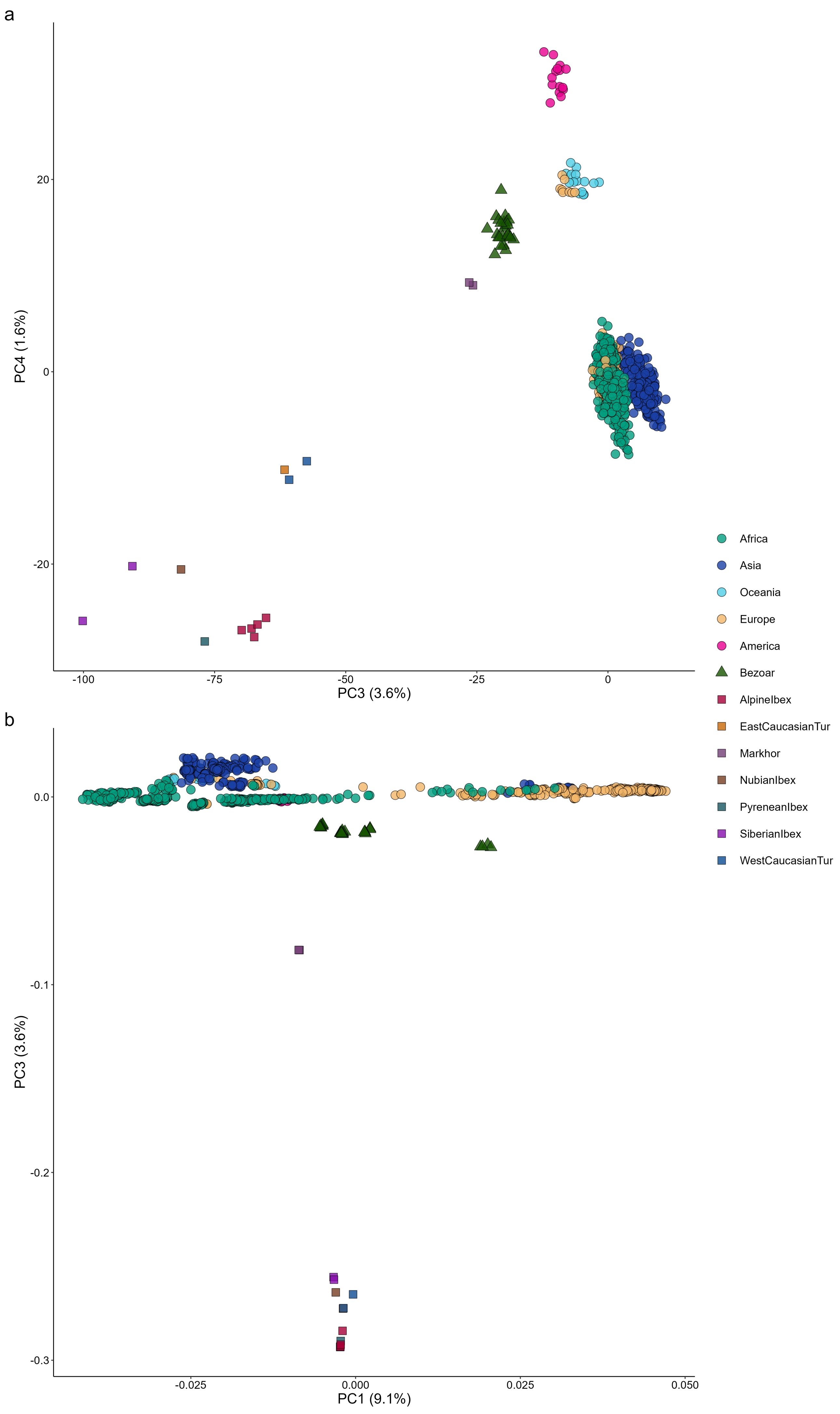

### Supplementary Figure S2

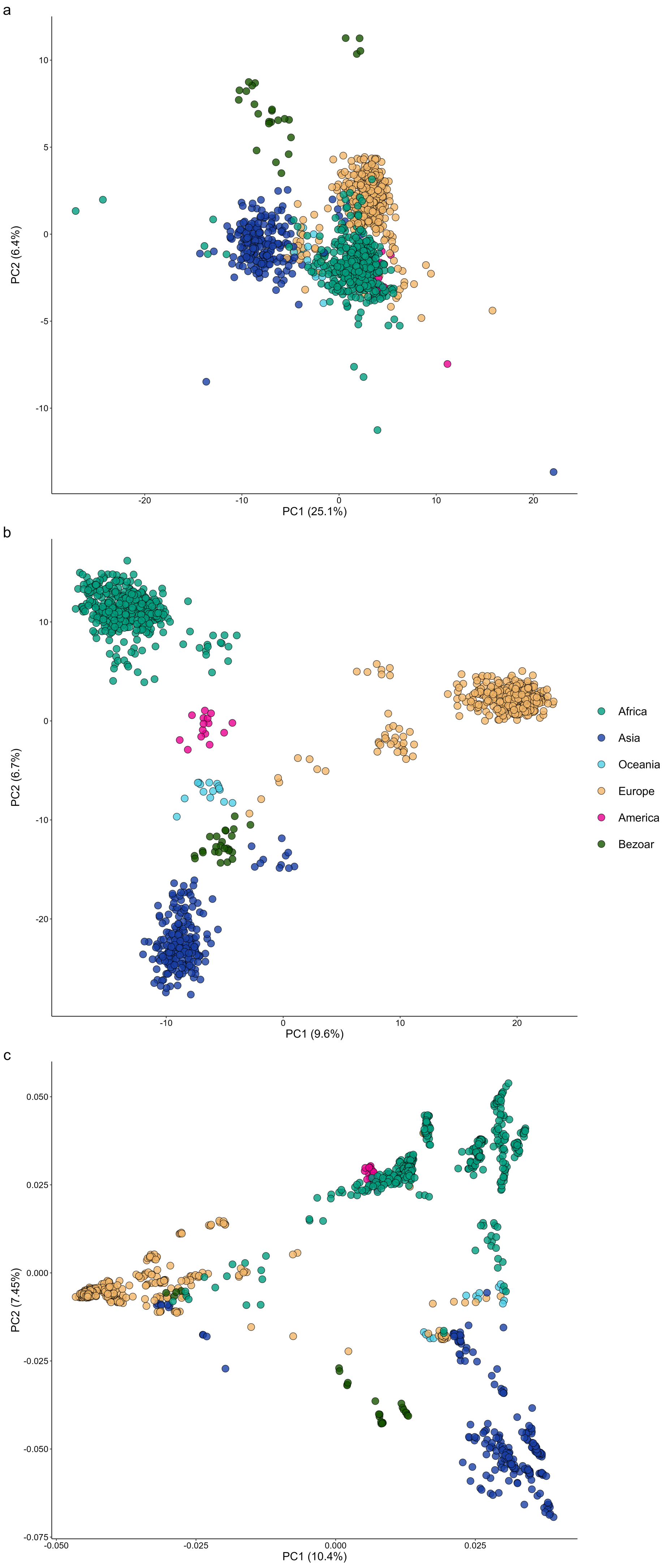

### Supplementary Figure S3

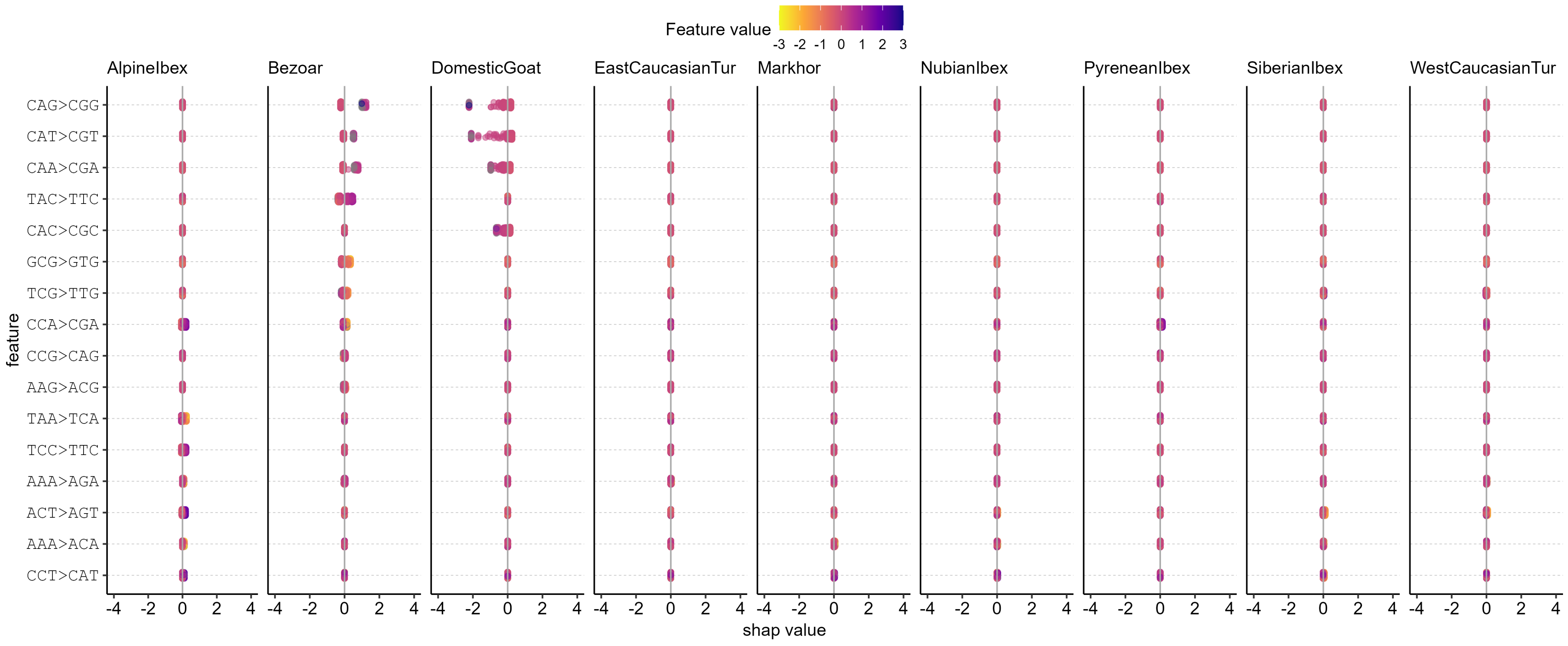

### Supplementary Figure S4

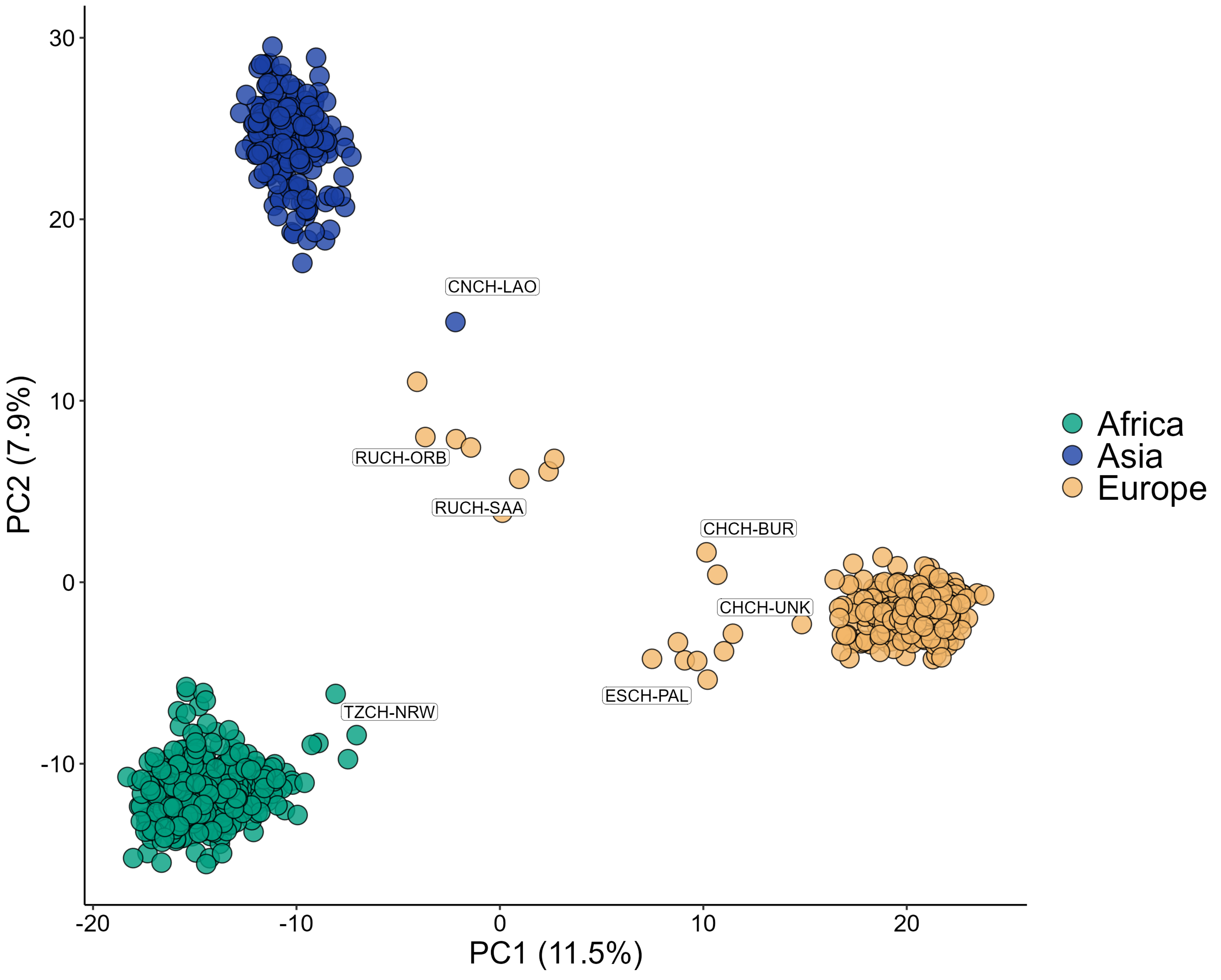

### Supplementary Figure S5

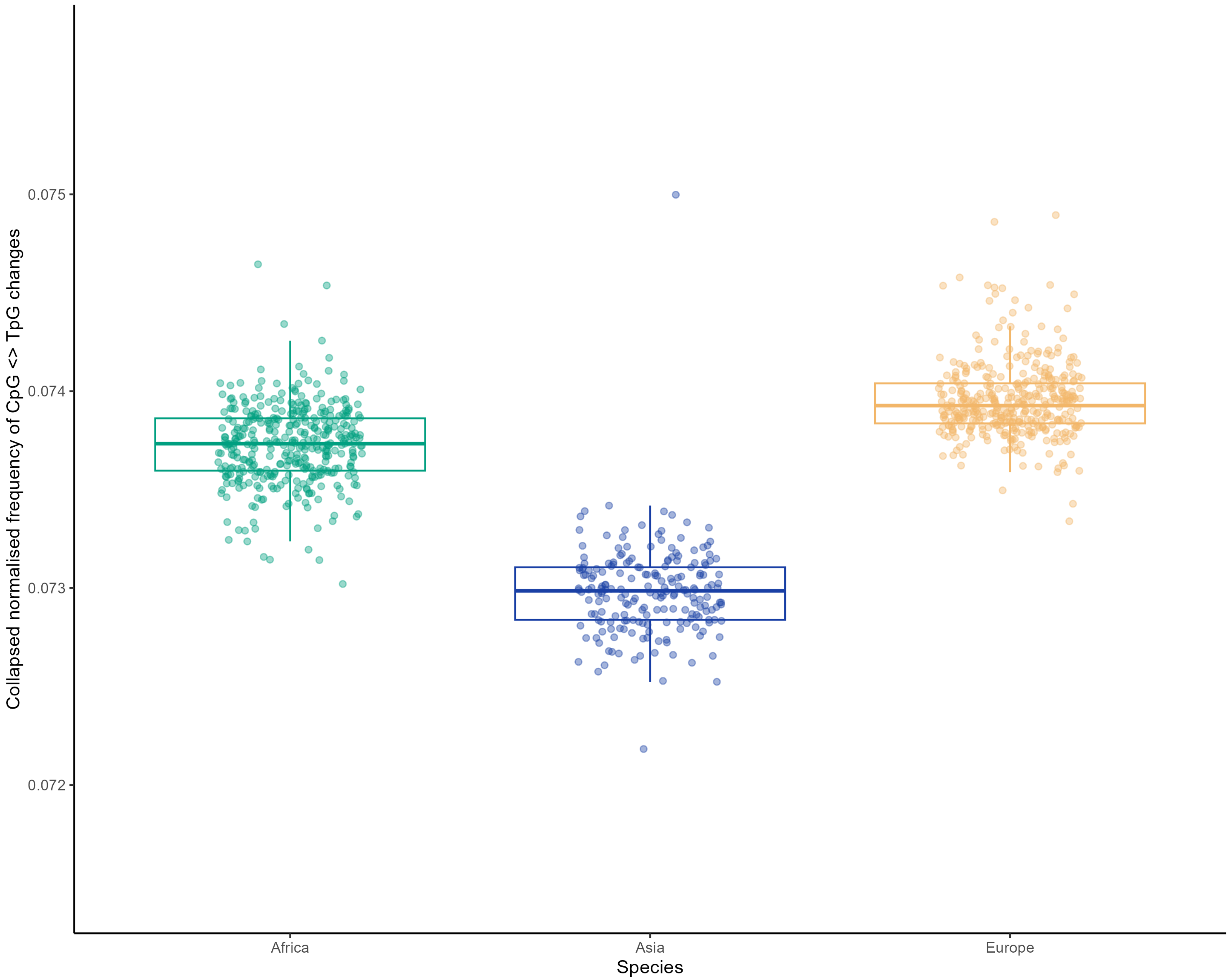

### Supplementary Figure S6

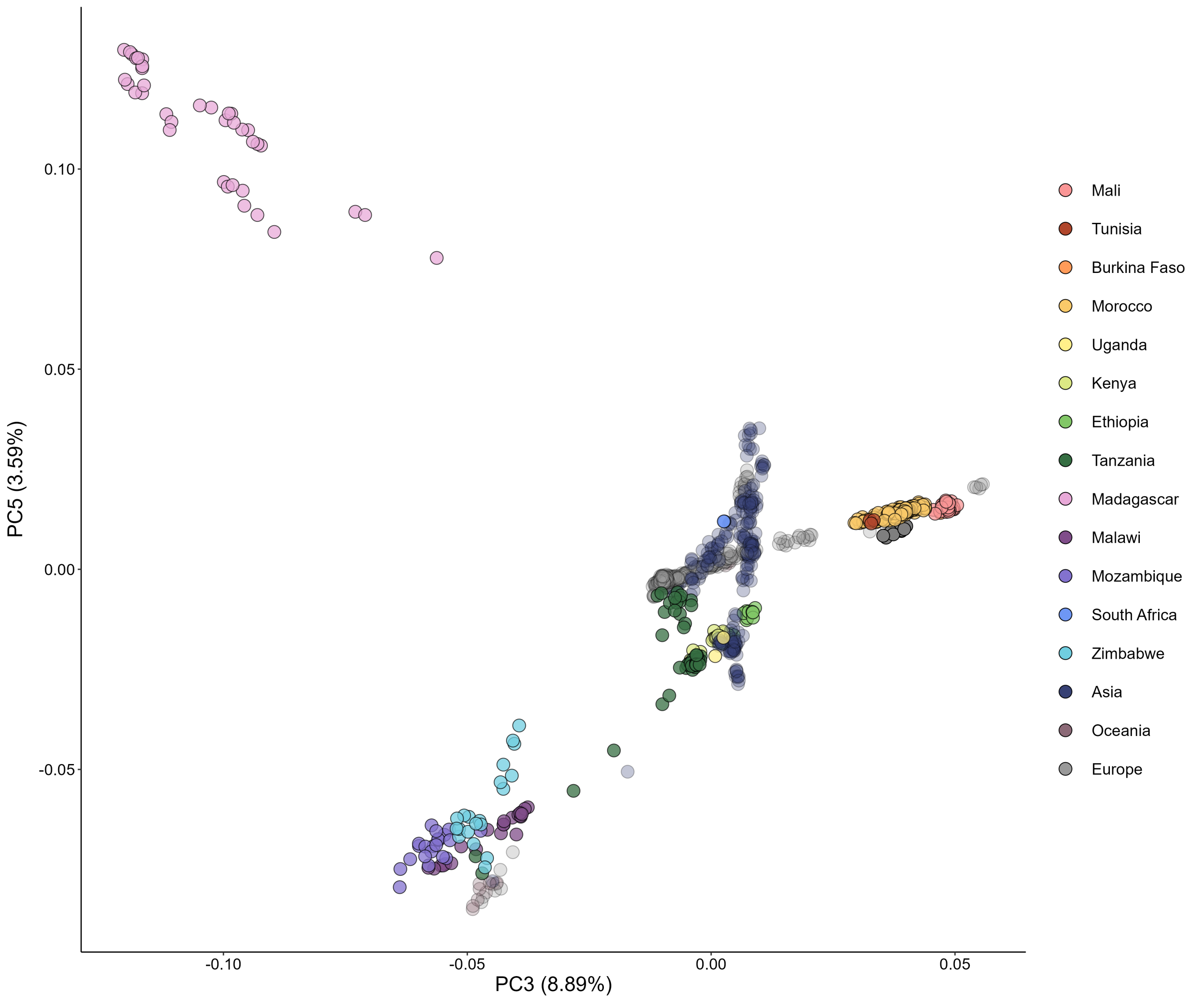

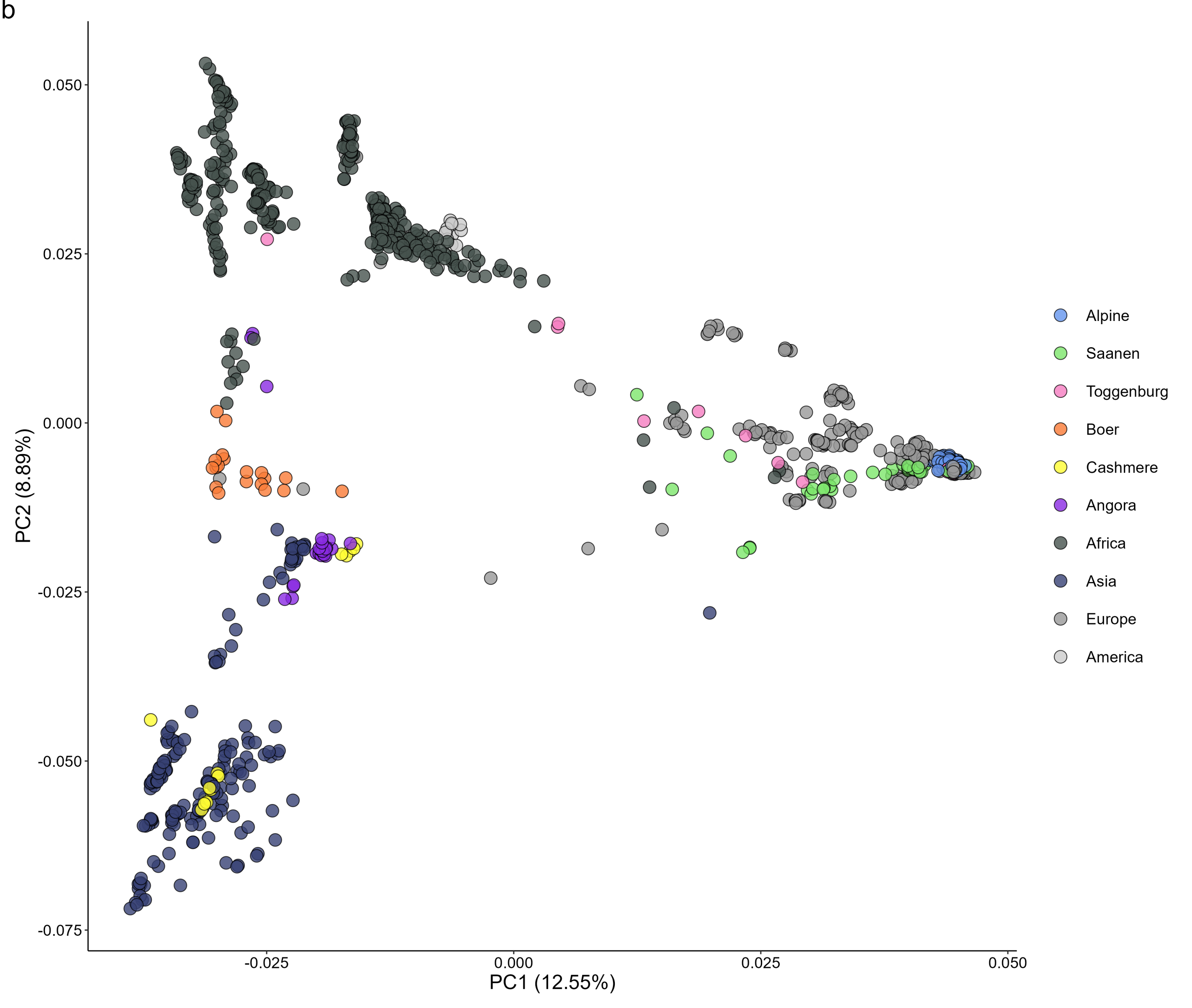

### Supplementary Figure S7

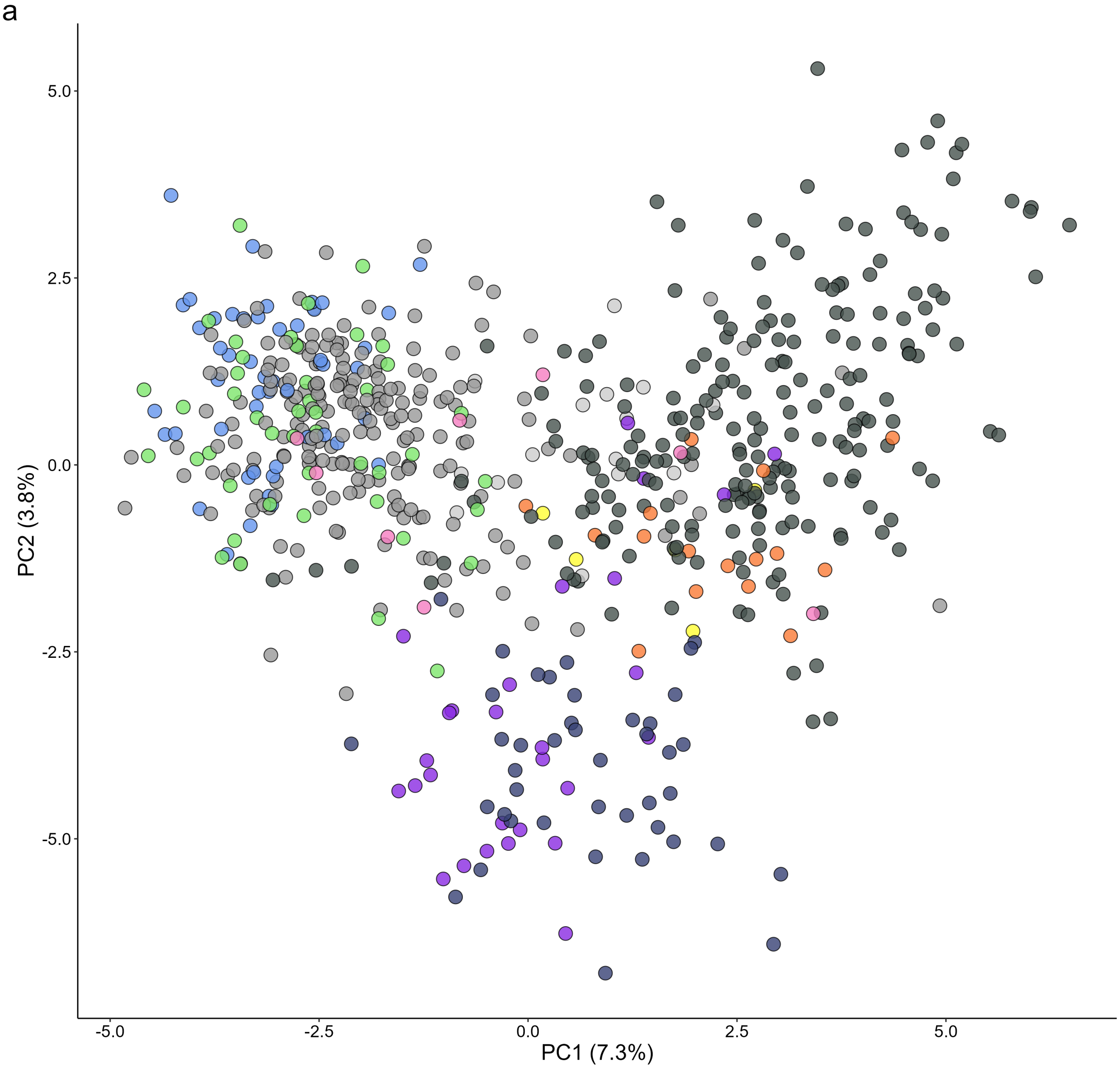

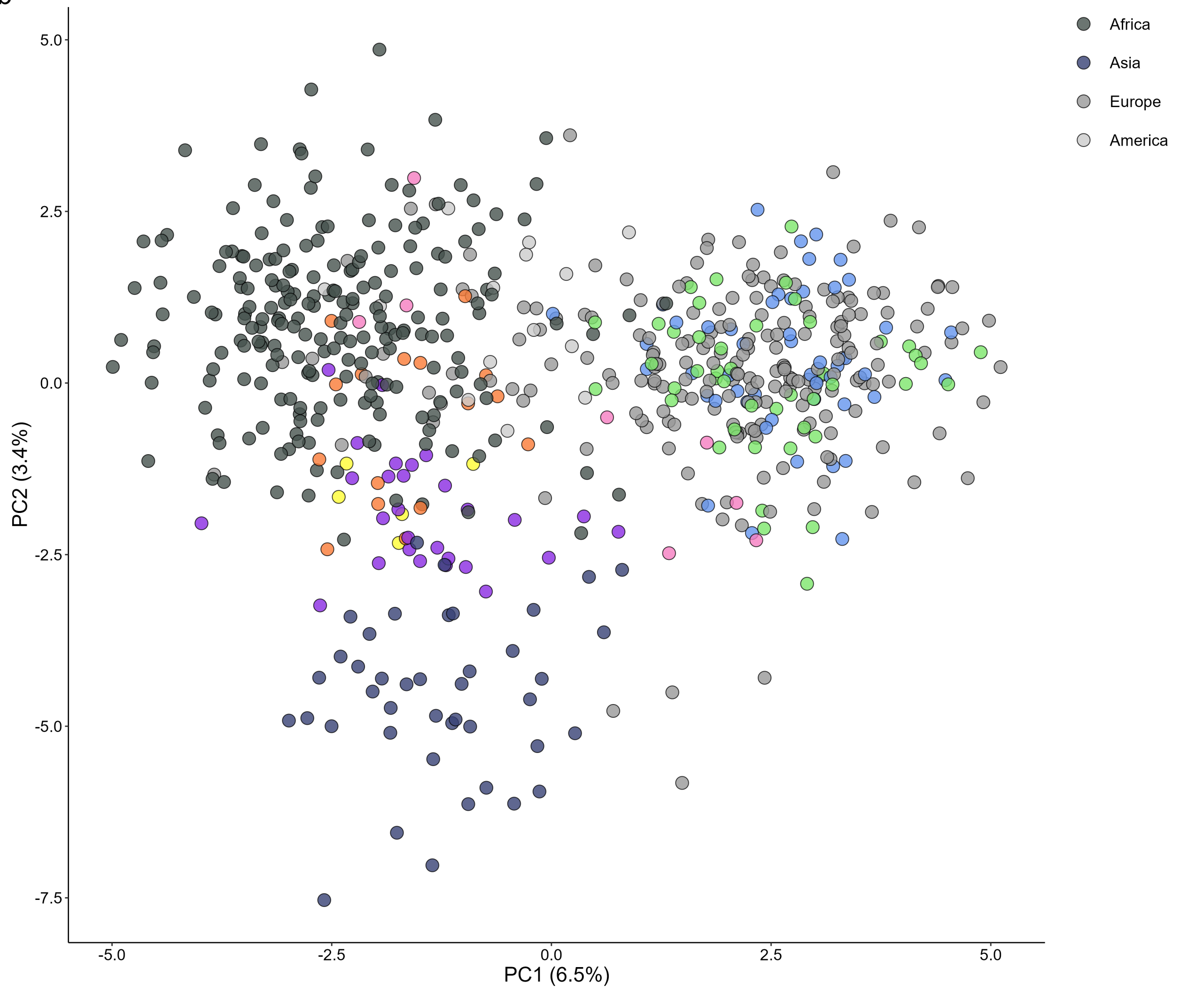
