## Appendix for "Linking geography and mutation profiles across goat species"

The VarGoats project members:

- Alessandra Crisà (Council for Agricultural Research and Economics CREA, Research Centre for Animal Production and Aquaculture, Monterotondo, Italy);
- Andrea Talenti (University of Glasgow, School of Biodiversity, One-Health and Veterinary Medicine, Glasgow, United Kingdom);
- Antonia Noce (Universitat Autònoma de Barcelona, CRAG Centre de Recerca en Agrigenòmica, Barcelona, Spain);
- Arianna Bionda (University of Milan, Dipartimento di Scienze Agrarie e Ambientali DiSAA, Milan, Italy);
- Arianna Manunza (Georg-August Universität, Fakultät für Forstwissenschaften und Waldökologie, Göttingen, Germany);
- Badr Benjelloun (INRAE-Moroc, Rabat, Morocco);
- Barbara Lazzari (National Research Council of Italy CNR, IBBA Istituto di Biologia e Biotecnologia Agraria, Milan, Italy; Università Cattolica del S. Cuore, Department of Animal Science,Food and Nutrition DIANA, Piacenza, Italy);
- Benjamin D. Rosen (USDA, Animal Genomics and Improvement Laboratory, Beltsville, MD 20705 USA);
- Bertrand Servin (INRAE, Génétique Physiologie et Systèmes d'Elevage GenPhySE, Castanet Tolosan, France);
- Carine Genet (INRAE, Génétique Physiologie et Systèmes d'Elevage GenPhySE, Castanet Tolosan, France);
- Carole Charlier (University of Liège, GIGA, Liège, Belgium);
- Caroline Leroux (Université Lyon 1, INRAE, Lyon, France);
- Coralie Danchin (IDELE, Génétique et gestion des populations animales, Paris, France);
- Cord Drögemüller (University of Bern, Institute of Genetics, Bern, Switzerland);
- Curt Van Tassell (USDA Animal Genomics and Improvement Laboratory, Beltsville, MD 20705 USA);
- Diego Alonso Vargas Donayre (Universitat Autònoma de Barcelona, CRAG Centre de Recerca en Agrigenòmica, Barcelona, Spain);
- Elena Petretto (Università Cattolica del S. Cuore, Department of Animal Science, Food and Nutrition DIANA, Piacenza, Italy);
- Emilio Mármol-Sánchez (Center for Evolutionary Hologenomics, The Globe Institute, University of Copenhagen, Copenhagen, Denmark);
- Emily Clark (EMBL-EBI European Bioinformatics Institute, Hinxton, United Kingdom);
- Emmanuelle Lerat (Université Lyon 1, CNRS Biometrie et Biologie Evolutive, Villeurbanne, France);

- François Pompanon (Université Grenoble Alpes, Laboratoire d'Ecologie Alpine, Grenoble, France);
- George Liu (USDA Animal Genomics and Improvement Laboratory, Beltsville, MD 20705 USA);
- Gwenola Tosser-Klopp (INRAE GenPhySE, Castanet Tolosan, France);
- Imen Baazaoui (Universitat Autònoma de Barcelona, CRAG Centre de Recerca en Agrigenòmica, Barcelona, Spain);
- Johannes Arjen Lenstra (Utrecht University, Faculty of Veterinary Medicine, Utrecht, Netherlands);
- Joram M. Mwacharo (International Centre for Agricultural Research in the Dry Areas ICARDA, Addis Ababa, Ethiopia);
- James Prendergast (University of Edinburgh, Roslin Institute, Midlothian, United Kingdom);
- Lin Jiang (Chinese Academy of Agricultural Sciences, Institute of Animal Science, Beijing, China);
- Jocelyn Turpin (Université Lyon 1, INRAE, Lyon, France);
- Jolijn Erven (University College Dublin, School of Agriculture and Food Science, Dublin, Ireland);
- Kevin Daly (University College Dublin, School of Agriculture and Food Science, Dublin, Ireland);
- Laura Botigué (Universitat Autònoma de Barcelona, CRAG Centre de Recerca en Agrigenòmica, Barcelona, Spain);
- Laurence Drouilhet (INRAE GenPhySE, Castanet Tolosan, France);
- Licia Colli (Università Cattolica del S. Cuore, Department of Animal Science, Food and Nutrition DIANA, Piacenza, Italy);
- Lingzhao Fang (University of Edinburgh, Roslin Institute, Midlothian, United Kingdom);
- Marina Naval (CSIRO Livestock genomics, St. Lucia, Queensland, Australia);
- Mahesh Neupane (USDA Animal Genomics and Improvement Laboratory, Beltsville, MD 20705 USA);
- Marcel Amills (Universitat Autònoma de Barcelona, CRAG Centre de Recerca en Agrigenòmica, Barcelona, Spain);
- Marco Milanesi (Università Cattolica del S. Cuore, Department of Animal Science, Food and Nutrition DIANA, Piacenza, Italy);
- Maria Gracia Luigi Sierra (Universitat Autònoma de Barcelona, CRAG Centre de Recerca en Agrigenòmica, Barcelona, Spain);
- Mario Barbato (Università degli Studi di Messina, Dipartimento di Scienze Veterinarie, Messina, Italy);
- Matteo Cortellari (University of Milan, Dipartimento di Scienze Agrarie e Ambientali DiSAA, Milan, Italy);

- Maxime Ben Braiek (INRAE, Génétique Physiologie et Systèmes d'Elevage GenPhySE, Castanet Tolosan, France);
- Mazdak Salavati (Scotland's Rural College SRUC, Dairy Research and Innovation Centre, Edinburgh, United Kingdom);
- Mingjing Wang (Universitat Autònoma de Barcelona, CRAG Centre de Recerca en Agrigenòmica, Barcelona, Spain);
- Paola Crepaldi (University of Milan, Dipartimento di Scienze Agrarie e Ambientali DiSAA, Milan, Italy);
- Philippe Bardou(INR AE, Génétique Physiologie et Systèmes d'Elevage GenPhySE, Castanet Tolosan, France);
- Pierre Faux (INRAE, Génétique Physiologie et Systèmes d'Elevage GenPhySE, Castanet Tolosan, France);
- Rachel Rupp (INRAE, Génétique Physiologie et Systèmes d'Elevage GenPhySE, Castanet Tolosan, France);
- Roberto Steri (Council for Agricultural Research and Economics CREA, Research Centre for Animal Production and Aquaculture, Monterotondo, Italy);
- Rudiger Brauning (AgResearch ltd., Bioeconomy Science Institute, Mosgiel, New Zealand);
- Alessandra Stella (National Research Council of Italy CNR, IBBA Istituto di Biologia e Biotecnologia Agraria, Milan, Italy);
- Stéphane Fabre (INRAE, Génétique Physiologie et Systèmes d'Elevage GenPhySE, Castanet Tolosan, France);
- Taina Figueiredo (Universitat Autònoma de Barcelona, CRAG Centre de Recerca en Agrigenòmica, Barcelona, Spain);
- Thomas Faraut (INRAE, Génétique Physiologie et Systèmes d'Elevage GenPhySE, Castanet Tolosan, France);
- Valentin Sorin (INRAE, GABI and GenPHYse, Jouy-en-Josas, France);
- Ke Wang (Universitat Autònoma de Barcelona, CRAG Centre de Recerca en Agrigenòmica, Barcelona, Spain);
- Clet Wandui Masiga (Tropical Institute of Development Innovations TRIDI, Kampala, Uganda);
- Wilson Nandolo (Lilongwe University of Agriculture and Natural Resources, Bunda Animal Breeding Centre, Lilongwe, Malawi);
- Yefang Li (Chinese Academy of Agricultural Sciences, Institute of Animal Science, Beijing, China);
- Zexi Cai (Aarhus university, Center for Quantitative Genetics and Genomics, Aarhus, Denmark).
