## Supplementary Materials for "Linking geography and mutation profiles across goat species"

**Supplementary Figure S1.** Principal components (PCs) 3 and 4 of SDM analysis (a) and PCs 1 and 3 of genotype analysis (b) showing the separation of the different wild *Capra* species.

**Supplementary Figure S2.** Principal component analysis showing the Bezoar's closeness to the Asian domestic goats using SNV (a), SDM (b), and genotypes (c).

**Supplementary Figure S3.** SHAP values showing the contribution of SNVs to the classification of *Capra* species. Positive SHAP values indicate variants that increase the probability of assignment to a given species, whereas negative values decrease it.

**Supplementary Figure S4.** SDM-based principal component analysis of UNREL_NOEXPORTED dataset, with intermediate groups named according to the Supplementary Table S1: CNCH-LAO = Laoshan breed from China; RUCH-ORB = Orenburg Chashmere breed from Russia; RUCH-SAA = Saanen breed from Russia; CHCH-BUR = Burlina breed from Switzerland; CHCH-UNK = Unknown breed from Switzerland; ESCH-PAL = Palmera breed from Spain (Canary Island); TZCH-NRW = Norwegian breed from Tanzania.

**Supplementary Figure S5.** Collapsed C>T changes in CpG context showing the depletion in the Asian goats.

**Supplementary Figure S6.** Principal component analysis of the genotypes, showing the separation of the Madagascar goats, similarly to the SDM principal component analysis (a), and the distribution of transboundary goats (b).

**Supplementary Figure S7.** Principal component analyses of the mutational profile computed from the first (a) and second (b) change composing the SDMs showing the distribution of transboundary goats.

**Supplementary Table S1.** List of samples and their belonging to the different datasets.

**Supplementary Table S2.** Results of the linear models for change types as a function of longitude, latitude, and their interaction terms, and correlations between each change type and geographic coordinates.
